# The RAM signaling pathway links morphology, thermotolerance, and CO_2_ tolerance in the global fungal pathogen *Cryptococcus neoformans*

**DOI:** 10.1101/2022.08.14.503895

**Authors:** Benjamin J. Chadwick, Tuyetnhu Pham, Xiaofeng Xie, Laura C. Ristow, Damian J. Krysan, Xiaorong Lin

## Abstract

The environmental pathogen *Cryptococcus neoformans* claims over 180,000 lives each year. Survival of this basidiomycete at host CO_2_ concentrations has only recently been considered an important virulence trait. Through screening gene knockout libraries constructed in a CO_2_-tolerant clinical strain, we found mutations leading to CO_2_ sensitivity are enriched in pathways activated by heat stress, including calcineurin, Ras1-Cdc24, cell wall integrity, and Regulator of Ace2 and Morphogenesis (RAM). Overexpression of Cbk1, the conserved terminal kinase of the RAM pathway, partially restored defects of these mutants at host CO_2_ or temperature levels. In ascomycetes such as *Saccharomyces cerevisiae* and *Candida albicans*, transcription factor Ace2 is an important target of Cbk1, activating genes responsible for cell separation. However, no Ace2 homolog or any downstream component of the RAM pathway has been identified in basidiomycetes. Through *in vitro* evolution and comparative genomics, we characterized mutations in suppressors of *cbk1*Δ in *C. neoformans* that partially rescued defects in CO_2_ tolerance, thermotolerance, and morphology. One suppressor is the RNA translation repressor Ssd1, which is highly conserved in ascomycetes and basidiomycetes. The other is a novel ribonuclease domain-containing protein, here named *PSC1*, which is present in basidiomycetes and humans but surprisingly absent in most ascomycetes. Loss of Ssd1 in *cbk1*Δ partially restored cryptococcal ability to survive and amplify in the inhalation and intravenous murine models of cryptococcosis. Our discoveries highlight the overlapping regulation of CO_2_ tolerance and thermotolerance, the essential role of the RAM pathway in cryptococcal adaptation to the host condition, and the potential importance of post-transcriptional control of virulence traits in this global pathogen.

## Introduction

There are over 278,000 cases of cryptococcal meningitis every year, causing over 180,000 deaths (Rajasingham et al., 2017). Cryptococcal meningitis is primarily caused by the ubiquitous environmental fungus *Cryptococcus neoformans*. Airborne spores or desiccated yeast cells of *C. neoformans* are inhaled into the lungs, where they are cleared or remain dormant until reactivation upon host immunosuppression (Casadevall & Perfect, 1998; Youbao Zhao, Lin, Fan, & Lin, 2019).

Litvintseva et al. found that most environmental *Cryptococcus* isolates cannot cause fatal disease in mouse models of cryptococcosis, despite having similar genotypes and *in vitro* phenotypes to known virulent isolates, including thermotolerance, melanization, and capsule production (Litvintseva & Mitchell, 2009). Mukaremera et al. also observed that *in vitro* phenotype assays for thermotolerance, capsule production, titan cell formation, or fluconazole heteroresistance, could not differentiate high-virulence strains from low-virulence strains (Mukaremera et al., 2019). These observations raise the possibility that other, unidentified virulence traits are important for *Cryptococcus* pathogenesis. Krysan and Lin laboratories demonstrated that tolerance to host levels of CO_2_ (∼5% CO_2_ in the host and ∼0.04% in ambient air) is likely a significant factor separating the potentially virulent natural isolates from the non-pathogenic environmental isolates that Litvintseva et al. tested (Krysan et al., 2019; Litvintseva & Mitchell, 2009).

The ability to adapt to host conditions is a prerequisite for cryptococcal pathogenesis. For instance, the ability of *C. neoformans* to replicate at human body temperature (≥37°C) has been extensively investigated. Many genes have been shown to be essential for thermotolerance (Perfect, 2006; Stempinski et al., 2021; Yang et al., 2017), including calcineurin which is currently being explored for antifungal drug development (Gobeil et al., 2021). By contrast, the underlying mechanisms or genes that play a role in CO_2_ tolerance have yet to be identified. Here, we set out to identify CO_2_-sensitive mutants and to gain the first insight into the genetic components involved in CO_2_ tolerance in *C. neoformans*.

## Results

### 1. CO_2_ sensitivity is independent of pH

Our previous work indicates that many *C. neoformans* environmental strains are sensitive to 5% CO_2_ when grown on buffered RPMI media, commonly used for mammalian cell cultures and testing antifungal susceptibility (Krysan et al., 2019). CO_2_ at host concentrations also acts synergistically with the commonly used antifungal drug fluconazole in inhibiting cryptococcal growth on buffered RPMI media. Because CO_2_ lowers the pH of aqueous environments, it is possible that the CO_2_ growth inhibitory effect or its synergy with fluconazole is simply due to lower medium pH. To address this question, we tested sensitivity to fluconazole of wild-type strain H99 using E-test on buffered RPMI media of either pH 6 or pH 7, with or without 5% CO_2_. In this E-test, the size of halo (clearance zone) reflects fungal susceptibility to fluconazole. As shown in Figure 1A, clearance zones were much larger in 5% CO_2_ relative to those in ambient air at both pH6 and pH7, indicating that CO_2_ sensitizes cryptococcal susceptibility to fluconazole. Furthermore, CO_2_ inhibits the growth of H99 at both pH 6 and pH 7 (smaller colony size in 5% CO_2_ relative to that in ambient air). Additionally, growth of CO_2_-sensitive environmental strain A7-35-23 (Krysan et al., 2019) was severely inhibited by 5% CO_2_ at both pH6 and pH7 (Figure 1B). In general, *C. neoformans* grows better at acidic pH (can grow well in pH3), and both A7-35-23 and H99 grew better at pH6 than at pH7 in ambient air (Figure 1B). Taken together, these results suggest that cryptococcal growth inhibition by CO_2_ is not simply due to lowered pH.

**Figure 1.**
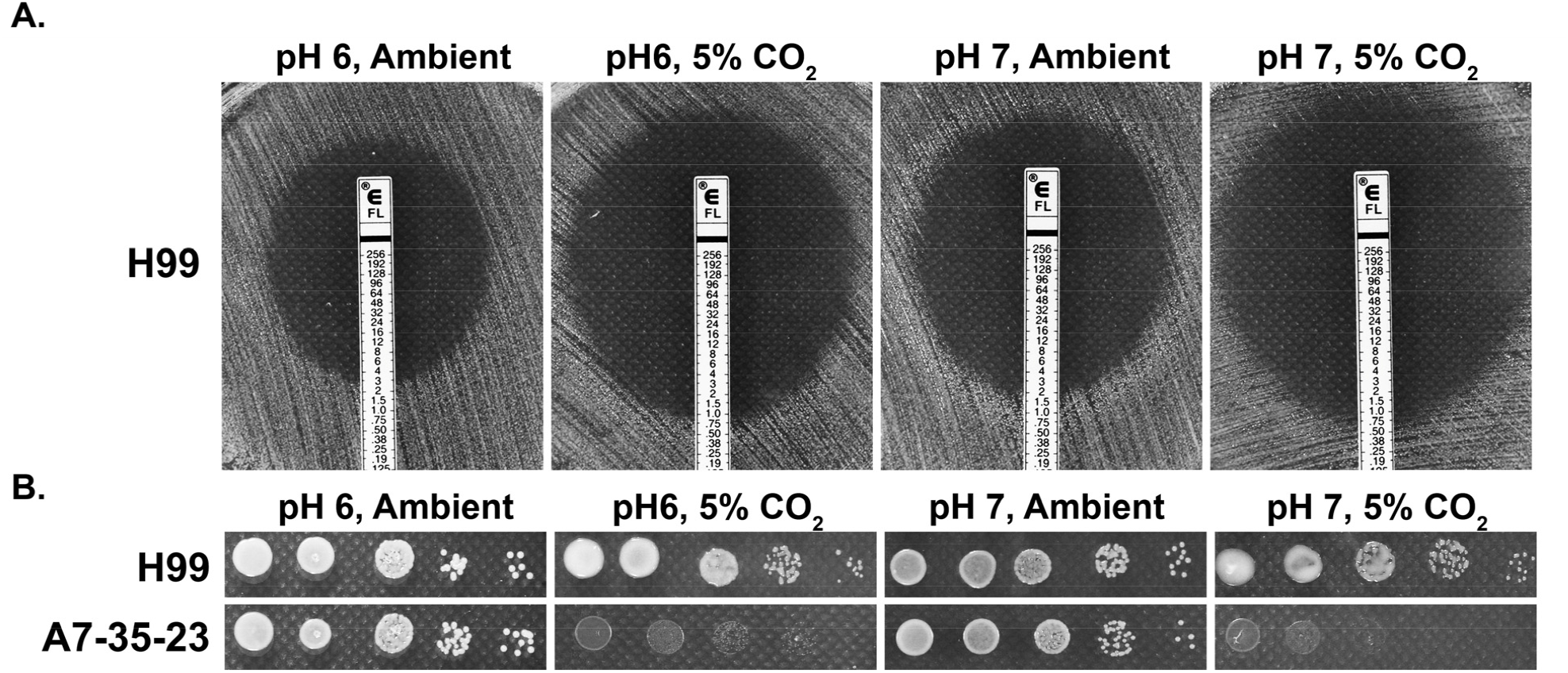
CO_2_ sensitivity is not simply due to lowered medium pH. (A) H99 cells were plated onto RPMI solid medium buffered to either pH 6 or pH 7. Fluconazole containing E-test strips were placed onto the lawn of H99 cells and the plates were incubated at 37°C in ambient air or in 5% CO_2_. The larger the halo surrounding the E-strip, the more sensitive the cells are to fluconazole. The intercept value of the halo with the E-strip is the minimal inhibitory concentration. (B) Cells of the previously identified CO_2_-tolerant H99 and CO_2_-sensitive A7-35-23 were serial diluted, spotted onto RPMI media buffered to pH 6 or pH7, and incubated at 37°C in ambient air or in 5% CO_2_.

### 2. Identifying genes important for CO_2_ tolerance

To identify genes involved in CO_2_ tolerance in *C. neoformans*, we screened gene deletion mutants constructed in the CO_2_-tolerant clinical reference strain H99. For large-scale screening, we were interested in using simplified medium. Accordingly, we tested the growth of two CO_2_-sensitive environmental strains and CO_2_-tolerant H99 in different levels of CO_2_ when cultured on the commonly used mycological YPD media. As expected, relative to H99, the CO_2_-sensitive strains A7-35-23 and A1-38-2 grew poorly at 5% CO_2_ and worse at 20% CO_2_ (Figure 2A). Using this approach, the following deletion mutant libraries were screened at 20% CO_2_ on YPD media: a set of strains previously constructed in our lab, the collections constructed by the Madhani lab, and a set generated in the Lodge Lab (Chun & Madhani, 2010). As some mutants are known to be temperature sensitive, we carried out the screens at 30°C rather than 37°C. Out of the over 5,000 gene knockout mutants screened, 96 were found to be sensitive to CO_2_ by visual observation (Table S1). We noticed that knockout mutants for multiple pathways known to be activated by heat stress are CO_2_ sensitive, including the Ras1-Cdc24 pathway, calcineurin, cell wall integrity (CWI), and <u>R</u>egulator of <u>A</u>ce2 and <u>M</u>orphogenesis (RAM). This finding indicates an overlapping nature of these two traits.

**Figure 2.**
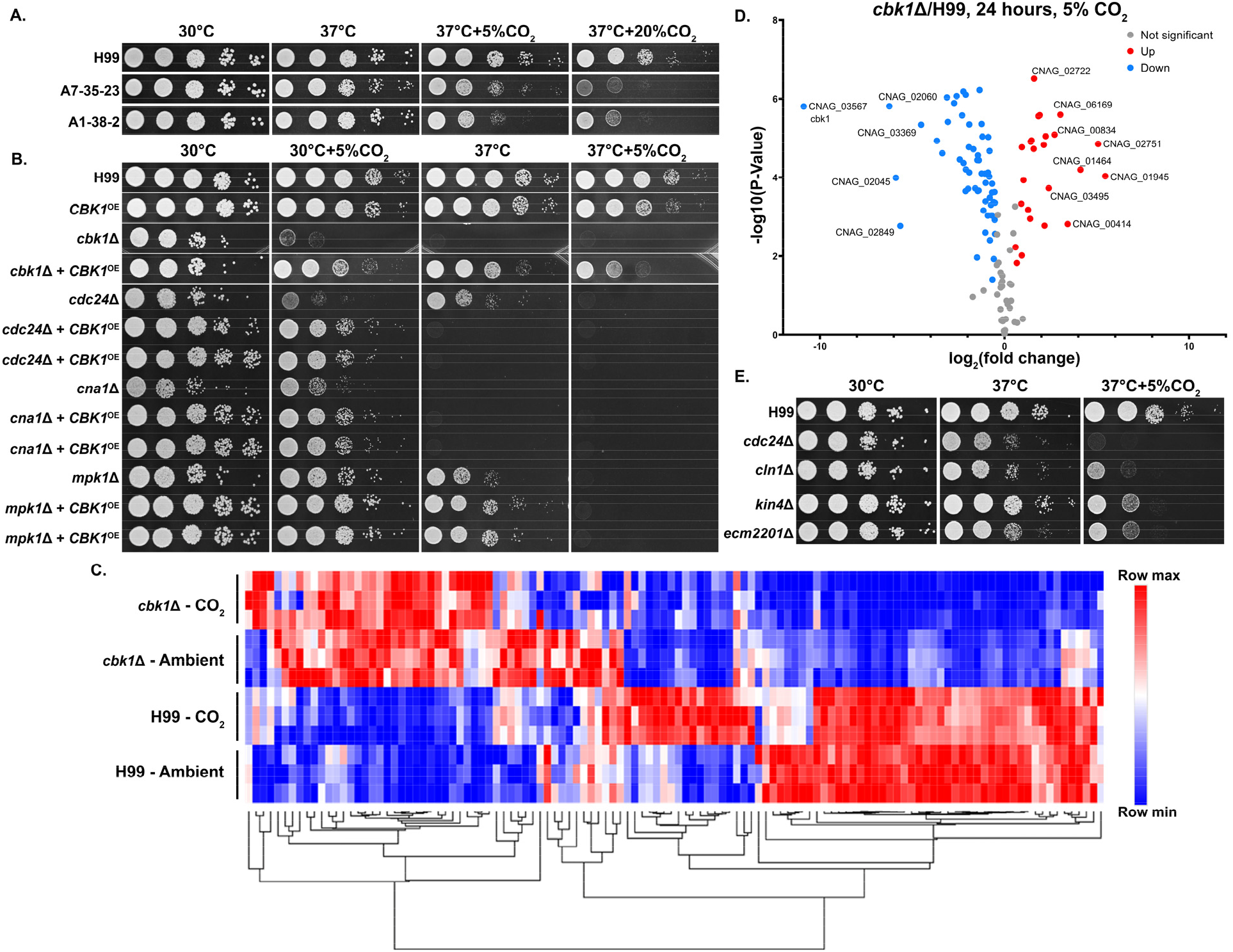
The RAM pathway effector kinase Cbk1 is critical for CO_2_ tolerance. (A). The clinical reference strain H99 and environmental strains A7-35-23 and A1-38-2 were grown overnight in YPD, serially diluted, and spotted onto solid YPD media plates. Photographs were taken two days after incubation in the indicated condition. (B) This serial dilution spotting assay was similarly performed for H99 and the mutants indicated. Two independent overexpression transformants for each mutant background were included as biological replicates. (C) Heatmap showing normalized total RNA counts of NanoString targets in H99 and *cbk1Δ* cultured at either ambient or 5% CO_2_, red indicates higher and blue indicates lower transcript abundance. (D) Volcano plot showing significantly differentially expressed transcripts (P value of <0.05) in the *cbk1Δ* compared to H99 in the 5% CO_2_ condition. (E) Serial dilution spotting assay of H99 and four of the mutants found in the deletion set screening to be CO_2_ sensitive which also correspond to significantly downregulated genes shown in the Volcano plot.

We were surprised by the absence of components of adenylyl cyclase-PKA pathway from the set of hits. In *Candida albicans*, the adenylyl cyclase pathway is crucial for the yeast-hypha transition in response to host levels of CO_2_ (Klengel et al., 2005). This pathway has also been proposed to play an important role for *Cryptococcus* to sense CO_2_ (Mogensen et al., 2006). However, we found that adenylyl cyclase pathway mutants showed no growth defects in CO_2_, including the adenylyl cyclase mutant *cac1*Δ, the adenylyl cyclase associated protein mutant *aca1*Δ, the alpha G protein subunit mutant *gpa1*Δ, and the cAMP-dependent protein kinase mutant *pkr1*Δ (*SI Appendix*, Figure S1). This indicates that growth defects in response to host levels of CO_2_ are likely independent of bicarbonate activation of adenylyl cyclase. This is not unexpected given that bicarbonate is not a limiting factor under the high level of CO_2_ used in our screen.

Because the calcineurin, Ras1-Cdc24, CWI, and RAM pathways are all activated at host temperature and were identified in our screen for CO_2_-sensitive mutants, we reasoned their downstream effectors may be related or genetically interact. As the RAM pathway effector kinase mutant *cbk1*Δ showed the most severe defect in thermotolerance and CO_2_ tolerance compared to the mutants of the other pathways, we first overexpressed the gene *CBK1* in the following mutants, *cdc24*Δ (Ras1-Cdc24), *mpk1*Δ (CWI), *cna1*Δ (Calcineurin), and the *cbk1*Δ mutant itself, and observed their growth at host temperature and host CO_2_ (Figure 2B). Overexpression was achieved by placing the *CBK1* open reading frame after the inducible *CTR4* promoter, which is highly activated in YPD media (Ory, Griffith, & Doering, 2004; Wang et al., 2014; Wang, Zhai, & Lin, 2012). The *CBK1* overexpression construct was specifically integrated into the “safe haven” locus *SH2* (Jianfeng Lin, Fan, & Lin, 2020; Upadhya et al., 2017) in each mutant strain background to avoid complications due to positional effect. As expected, the growth defect of the *cbk1*Δ mutant at 37°C with and without 5% CO_2_ were largely restored by *CBK1* overexpression. At 30°C, overexpression of *CBK1* restored the growth of the *mpk1*Δ mutant, the *cna1*Δ mutant, and the *cdc24*Δ mutant in the CO_2_ condition. In terms of thermotolerance, overexpression of *CBK1* restored growth of *mpk1*Δ but not *cna1*Δ, while the growth defect of *cdc24*Δ at 37°C was exacerbated. *CBK1* overexpression failed to rescue growth of any of these mutants when both stressors were present (37°C + 5% CO_2_). We found that overexpression of *CBK1* in the WT H99 background caused a modest growth defect at 37°C + 5% CO_2_. Thus, the detrimental effects from *CBK1* overexpression under this growth condition may partially explain its inability to fully rescue growth of these tested CO_2_-sensitive mutants. The reciprocal overexpression of *CDC24, MPK1*, or *CNA1* in the *cbk1*Δ mutant background did not restore growth under 37°C and/or 5% CO_2_ (*SI Appendix*, Figure S2). These results support a hypothesis that Cbk1 integrates multiple stress response pathways to regulate both CO_2_ tolerance and thermotolerance.

To determine the extent of Cbk1’s role in CO_2_ tolerance, we conducted NanoString gene expression profiling of the WT H99 and *cbk1*Δ mutant cultured in ambient air and in 5% CO_2_ at 30°C (Figure 2C). Transcript levels of 118 gene was measured and those genes were chosen based on RNA sequencing results from a separate study. In that study, these genes were differentially expressed in CO_2_ vs ambient air conditions in either two CO_2_-sensitive or two CO_2_-tolerant natural strains (Dataset S1). Out of these 118 CO_2_-associated genes, 81 were found to be significantly differentially expressed in the *cbk1*Δ mutant in both ambient air and in 5% CO_2_, indicating they are intrinsically dysregulated in the *cbk1*Δ mutant.

57/81 of these genes are downregulated and 24/81 upregulated compared to the WT H99 strain (Figure 2D). Interestingly, 16/57 of the downregulated genes were also hits in our deletion set screening. We confirmed sensitivity to host CO_2_ conditions for four of these mutants by spotting assay (Figure 2E).

Taken together, this transcriptomic profiling shows that loss of Cbk1 significantly affects the expression of CO_2_-related genes.

### 3. The RAM signaling pathway is critical for normal morphology, thermotolerance, and CO_2_ tolerance

The RAM pathway effector kinase Cbk1 is part of the NDR/LATS family of kinases, which is conserved from yeast to humans and affects a wide range of cellular functions including cell-cycle regulation. Through our genetic screen for CO_2_-sensitive mutants, we found that all tested *Cryptococcus* RAM pathway mutants are extremely sensitive to 5% CO_2_ and high temperature, and they show no growth at 37°C + 5% CO_2_ (Figure 3A). In ascomycetes such as *S. cerevisiae* and *C. albicans*, RAM pathway mutants are defective in cytokinesis and exhibit loss of polarity, resulting in enlarged round cells that cluster together (Saputo, Chabrier-Rosello, Luca, Kumar, & Krysan, 2012)(Figure S3). In contrast, though defective in cytokinesis (Walton, Heitman, & Idnurm, 2006), *Cryptococcus* RAM pathway mutants are hyper-polarized and constitutively form clusters of elongated pseudohyphal cells (Figure 3B). Moreover, we found that while the *C. albicans* homozygous *cbk1*ΔΔ mutant exhibits a general growth defect compared to the wild-type control, it shows no apparent specific growth defect at 37°C with or without 5% CO_2_ (Figure S3). These results suggest that, although the RAM pathway is conserved in its role in cytokinesis, the effects of its downstream targets are divergent between ascomycetes and basidiomycetes.

**Figure 3.**
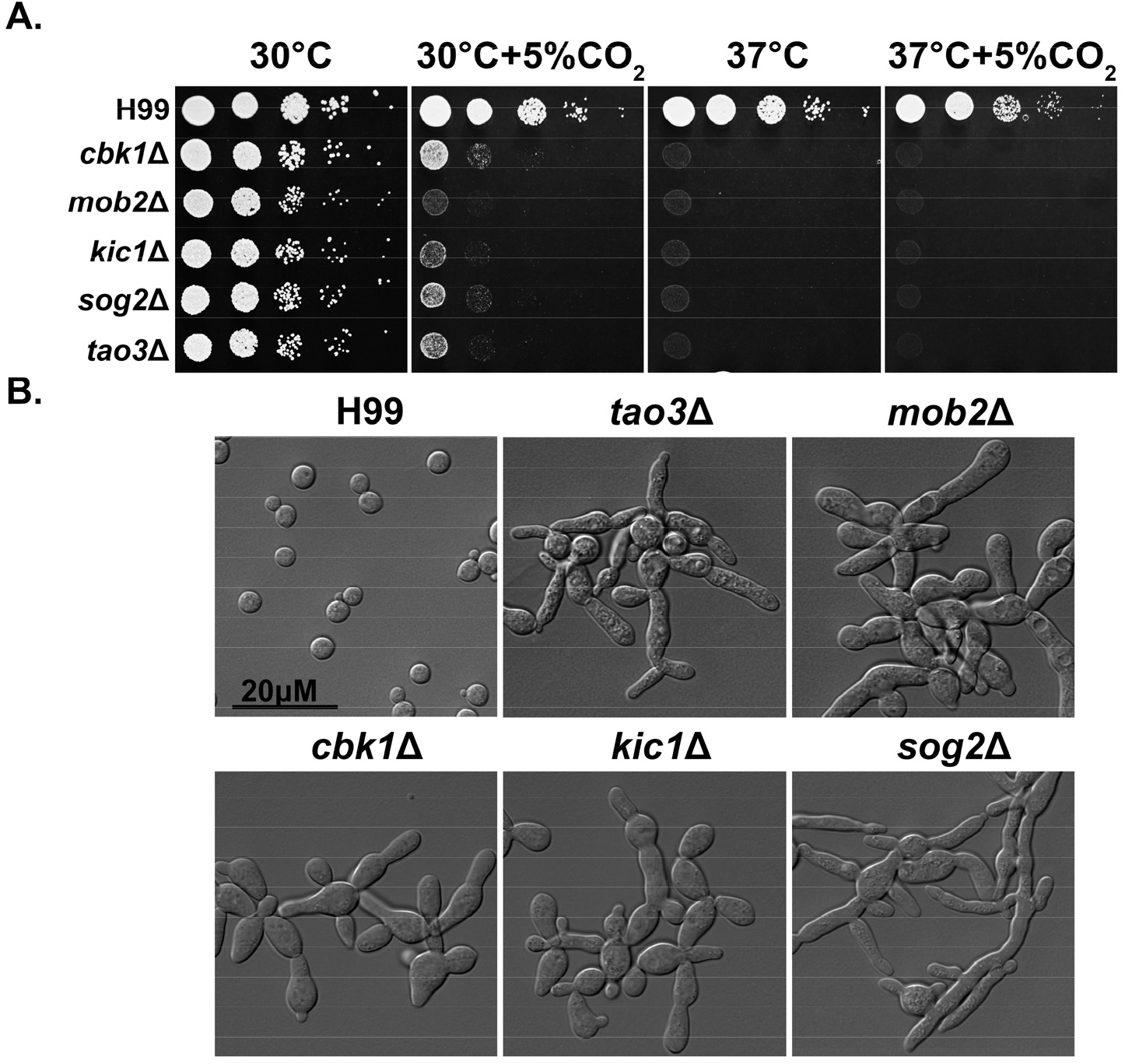
The RAM pathway is critical for normal morphology, thermotolerance, and CO_2_ tolerance. (A) *C. neoformans* WT H99 and RAM pathway mutants were serially diluted, spotted onto YPD medium, and incubated for two days at the indicated condition. (B) The cellular morphology of *Cryptococcus neoformans* WT H99 and RAM pathway mutants cultured in YPD medium.

### 4. Suppressors of the *cbk1*Δ mutant show improved growth at host conditions

In ascomycetes, Ace2 is a key downstream transcription factor of the RAM pathway (hence in the name of RAM − regulator of <u>A</u>ce2 and <u>m</u>orphogenesis), which is important for the activation of genes responsible for cell separation. However, no homolog to Ace2 has been identified in *Cryptococcus* or other basidiomycetes. Furthermore, no downstream targets of the RAM pathway have been identified in any basidiomycetes. To investigate potential downstream effectors of the RAM pathway in *Cryptococcus*, we screened for spontaneous suppressor mutants of *cbk1*Δ. To do so, *cbk1*Δ mutant cells from an overnight culture in liquid YPD at 30°C were plated onto solid YPD media and incubated for two days at 37°C + 5% CO_2_. Out of >1×10^8^ cells plated and cultured under this condition that is inhibitory for growth of the original *cbk1*Δ mutant, 11 suppressor colonies were isolated for further examination and sequencing. All the suppressor isolates showed dramatically improved growth over the original *cbk1*Δ mutant at 37°C and modestly improved growth at 37°C + 5% CO_2_ (Figure 4C). Based on their distinctive phenotypes, the 11 suppressors were classified into two groups: *sup1* (2/11) and *sup2* (9/11). Shorter chains of cells in both groups indicate a partial restoration in cytokinesis (Figure 4D). The *sup2* group has slightly improved growth at 37°C + 5% CO_2_ and forms shorter chains of cells compared to the *sup1* group (Figure 4C, 4D). Besides these observations, *sup1* and *sup2* displayed similar phenotypes in all other cryptococcal virulence traits tested, including melanin production, capsule, urease activity, and cell wall stress tolerance (*SI Appendix*, Figure S4).

**Figure 4.**
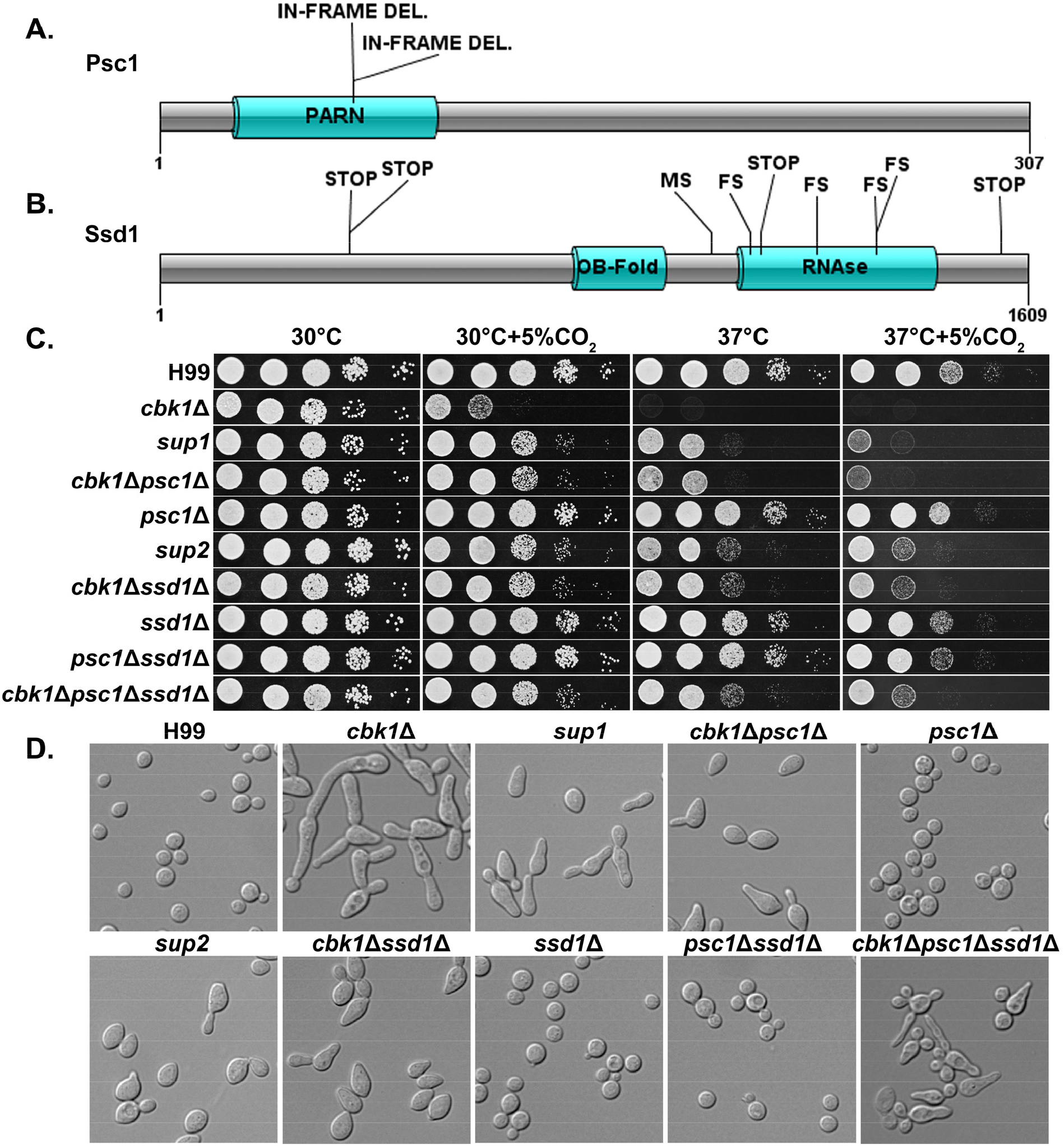
Natural suppressors of the RAM pathway *cbk1*Δ mutant restore multiple defects. (A) Protein diagram of Psc1 showing the effects and positions of suppressor mutations in the two *sup1* type natural suppressors. (B) Protein diagram of Ssd1 and the effects and positions of suppressor mutations in Ssd1 in the nine *sup2* type natural suppressors. STOP indicates a nonsense mutation, MS a missense mutation, and FS a frameshift mutation. Asterisks (*) indicate mutations in the same suppressor strain. (C) Serial dilutions of H99 and the mutant strains were spotted onto YPD agar media and incubated for two days in the indicated condition to observe growth. (D) The cellular morphology of H99 and the mutant strains in liquid YPD cultures were examined under microscope.

Along with the original *cbk1*Δ mutant, we sequenced the genomes of the 11 *cbk1*Δ suppressors. By comparing their genome sequences with each other and with the original *cbk1*Δ mutant, we found that both *sup1* type suppressor mutants contained a disruptive in-frame deletion at the same location in *CNAG_01919*, which encodes a putative Poly(A)-specific ribonuclease (PARN) domain-containing protein (Figure 4A). Interestingly, while this domain is found in *S. pombe* (Marasovic, Zocco, & Halic, 2013), we did not identify this domain in the genomes of *S. cerevisiae, C. albicans* or other ascomycetes. By contrast, the PARN domain is common in basidiomycetes and higher eukaryotes. The in-frame deletion results in a change of two amino acids within the predicted PARN domain, the only discernable domain present in this protein. We named this previously uncharacterized gene <u>P</u>artial <u>S</u>uppressor of *<u>c</u>bk1*Δ (*PSC1*). All of the 9 *sup2* isolates contained loss of function or missense mutations in the gene *CNAG_03345* (Figure 4B), which encodes an RNA-binding protein homologous to *S. cerevisiae* Ssd1p, a known suppressor of *cbk1*Δ phenotypes in the model budding yeast. In *S. cerevisiae*, Ssd1p represses transcript translation and is negatively regulated by Cbk1p phosphorylation (Jansen, Wanless, Seidel, & Weiss, 2009; Wanless, Lin, & Weiss, 2014).

To confirm that the mutations in *SSD1* and *PSC1* are responsible for suppressing *cbk1*Δ, we created *cbk1*Δ*ssd1*Δ and *cbk1*Δ*psc1*Δ double mutants together with the control single mutants *ssd1*Δ and *psc1*Δ. Indeed, relative to *cbk1*Δ, the double mutants showed reduced sensitivity to host temperature and CO_2_ levels (Figure 4C), similar to the natural suppressor mutants. Likewise, the morphology of the double mutants resembles that of the spontaneous suppressor mutants (Figure 4D). The deletion of *SSD1* and *PSC1* alone in the wild-type background did not yield any discernable phenotype. The results confirm that loss of function in Ssd1 and Psc1 is responsible for the restoration of the *cbk1*Δ mutant’s growth defects observed in the natural suppressors. Interestingly, *sup2* and the *cbk1*Δ*ssd1*Δ mutants both grew noticeably better than *sup1* and *cbk1*Δ*psc1*Δ at 37°C and 37°C + 5% CO_2_. To test the genetic interaction between the two suppressor genes *SSD1* and *PSC1*, we created a triple *cbk1*Δ*psc1*Δ*ssd1*Δ mutant and the control strain *psc1*Δ*ssd1*Δ. The *psc1*Δ*ssd1*Δ control strain did not exhibit any defect and grew similarly well to either single mutant or the wild type (Figure 4C). The triple mutant *cbk1*Δ*psc1*Δ*ssd1*Δ grew similarly well as *sup2* or *cbk1*Δ*ssd1*Δ at 37°C + 5% CO_2_ (Figure 4C). However, the triple mutant displayed aberrant morphology and budding defects which are not observed in the natural suppressor mutants or the *cbk1*Δ*ssd1*Δ and *cbk1*Δ*psc1*Δ double mutants (Figure 4D). These results suggest that Psc1 and Ssd1 may function in the same pathway in regulating thermotolerance and CO_2_ tolerance, but their downstream effects on cell separation may be overlapping but distinct.

To determine if the suppressor mutations restore transcript abundance of the differentially expressed genes under CO_2_ in *cbk1*Δ, we compared the profiles of *cbk1*Δ to the two suppressor mutants: *sup1* and *sup2*. Overall, we found that the spontaneous suppressors do not restore transcript abundances of most differentially expressed genes in *cbk1*Δ to WT levels (*SI Appendix*, Figure S5), suggesting that post-transcriptional regulation might play a role in CO_2_ tolerance.

### 5. Spontaneous suppressors of *cbk1*Δ mutant show improved ability to survive and amplify in the host

RAM mutants have previously been found to be attenuated in virulence in the invertebrate wax moth larva infection model and mouse intranasal infection models. Occasionally, cryptococcal strains with point mutations in RAM genes cause death of mice when revertant mutations occur (Magditch, Liu, Xue, & Idnurm, 2012). Consistently, we found that the *cbk1*Δ mutant shows severe defect in growth at host temperature and CO_2_ levels. Because *sup1* and *sup2* both largely restored growth at 37°C but only modestly restored growth at 37°C + 5% CO_2_, we decided to test if and how much these suppressor mutations could restore the virulence defect of *cbk1*Δ. We infected mice with 1 × 10^4^ cells of WT, *cbk1*Δ, *sup1*, or *sup2* intranasally. In this intranasal infection model, the WT H99 strain establishes lung infection first and typically disseminates to other organs including the brain at about 7-10 days post-infection (DPI). Mice infected by H99 normally reach clinical endpoint around 3-4 weeks post-infection and these mice have a high fungal burden in the lungs, brain, and kidney (Chadwick & Lin, 2020; Jianfeng Lin et al., 2022). As expected, all mice infected by H99 were moribund by DPI 26 (Figure 5A) while the *cbk1*Δ mutant failed to cause any mortality when we terminated the experiment at DPI 60. Surprisingly, *sup1* and *sup2* strains did not cause any mortality either. The organ fungal burden, however, revealed differences in virulence levels between these strains. At the time of euthanasia for H99-infected mice (prior to DPI 26), the median fungal burden in the lungs, brains, and kidneys was 2.1×10^8^, 1.4×10^6^, and 2.4×10^4^ CFUs per organ respectively (Figure 5B). As expected, mice completely cleared the *cbk1*Δ cells when examined at DPI 35. Surprisingly, despite largely restored growth at 37°C, *sup1* was completely cleared from the mouse lungs by DPI 35, similar to the *cbk1*Δ mutant. In comparison, although *sup2* did not cause any death during the study period, it was able to grow in mouse lungs and the lung CFUs were ∼10-fold higher than the original inoculum when examined at DPI 35. The *sup2* strain maintained the same high fungal cell count in the lungs even at DPI 60 (Figure 5B), indicating that it can persist in the animals for a long time. The only obvious *in vitro* difference observed between *sup1* and *sup2* was better growth of *sup2* at host CO_2_ levels, which may explain the difference in their ability to propagate and persist in the mouse lungs.

**Figure 5.**
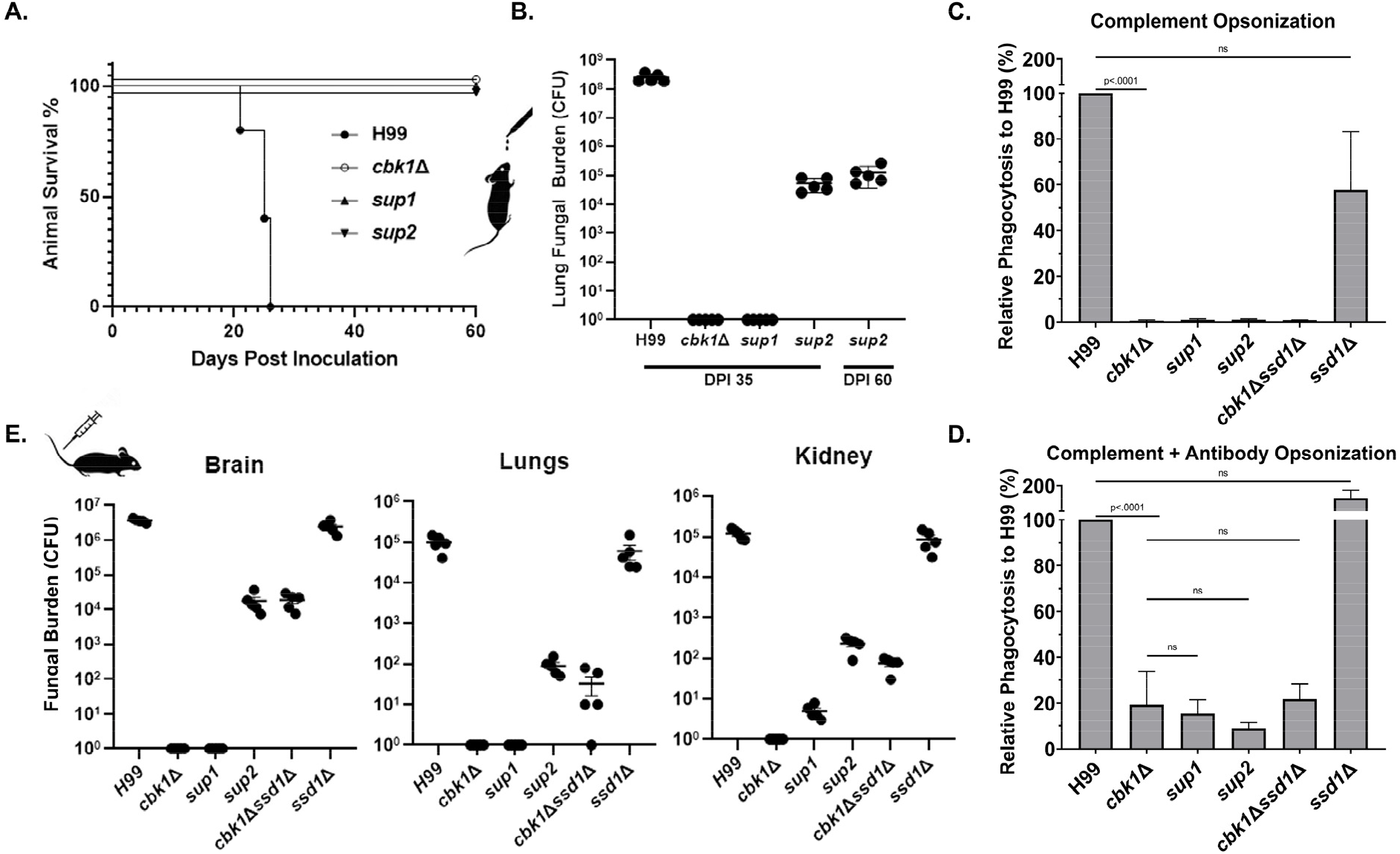
Suppressor mutants are partially restored for phagocytosis and can disseminate in the intravenous infection model of cryptococcosis. (A) Mice were infected with 1×10^4^ fungal cells intranasally, and their survival was monitored for 60 days post infection. (B) At day 35 post infection (DPI 35) and at the time of termination (DPI 60), 5 out of 10 mice per group for the *cbk1*Δ mutant, *sup1* and *sup2* groups were harvested for brains, kidneys, and lungs. For H99 infected mice, they were euthanized at their clinical end point (all before DPI 26). Tissue homogenate was serially diluted and plated onto YNB medium to count the colony forming units (CFUs) to measure the fungal burden per organ. (C) Murine macrophage J774A.1 cells were co-incubated with 2×10^6^ cryptococcal cells opsonized with serum from naïve mouse for two hours. Nonadherent or phagocytosed cells were washed, and cryptococcal cells were released and then serially diluted before plating onto YNB medium for measurement of colony forming units (CFUs). Statistical analysis was performed on the percentage of phagocytosed cells relative to H99, using one-way ANOVA. A p-value < 0.05 is considered statistically significant. (D) The same as above, except opsonization was performed with serum of mice vaccinated against cryptococcosis. (E) Mice were challenged with 1×10^5^ cryptococcal cells intravenously. At day 5 post infection, five mice per group were sacrificed. Brains, kidneys, and lungs of euthanized mice were dissected and homogenized. Serial dilutions were plated to count CFUs for quantification of fungal burden per organ.

Although the spontaneous suppressor *sup2* was able to grow and survive in the mouse lungs, there was no fungal burden detected in the brain or the kidney at DPI 35 or 60 (all zeros in all mice examined), indicating a failure in dissemination. We considered two hypotheses: 1) Inability of suppressor *sup2* to escape from the lungs; 2) Inability of suppressor *sup2* to penetrate other organs from the blood. Because one way that *C. neoformans* disseminates from the lungs to other organs is by a “Trojan Horse” mechanism, where *Cryptococcus* travels within the mobile host phagocytes (Kechichian, Shea, & Del Poeta, 2007; Santiago-Tirado, Onken, Cooper, Klein, & Doering, 2017), we examined phagocytosis of the *cbk1*Δ mutant and its suppressors to test the first hypothesis. We expect that cryptococcal mutants defective in being phagocytosed by host cells might be defective in dissemination, and the *cbk1*Δ mutant was previously found to have poor phagocytosis index (J. Lin, Idnurm, & Lin, 2015). Here, we co-cultured murine macrophage JA774 cells with H99, *cbk1*Δ, *sup1, sup2*, the double mutant *cbk1*Δ*ssd1*Δ, or the control single mutant *ssd1*Δ. Opsonization was performed using either naïve mouse serum (complement mediated phagocytosis) or serum from mice vaccinated against cryptococcosis (complement + antibody mediated phagocytosis)(Jianfeng Lin et al., 2022; Zhai et al., 2015). Consistent with our previous finding, phagocytosis of *cbk1*Δ was extremely low (∼1% of the WT H99 level under complement mediated phagocytosis, Figure 5C). Opsonization with serum from vaccinated mice increased phagocytosis of *cbk1*Δ and the suppressor mutants, but the phagocytosis indexes of these mutants were still only 20% or less than that of the wildtype (Figure 5D). In both phagocytosis experiments, the suppressor mutants did not show rescued phagocytosis. The poor phagocytosis of the *cbk1*Δ mutant and its suppressors may have contributed to their lack of dissemination from the lungs to the other organs in the inhalation infection mouse model of cryptococcosis.

To test the second hypothesis, we infected mice intravenously with H99, *cbk1*Δ, *sup1, sup2*, the double mutant *cbk1*Δ*ssd1*Δ, or the control single mutant *ssd1*Δ. In this intravenous infection model, the barrier of the lungs is bypassed. H99 cells are expected to disseminate to the brain and other organs within hours. Because of H99’s rapid dissemination in this model, infected mice typically reach clinical endpoint after 1 week. Therefore, we euthanized mice at DPI 5 before H99-infected mice would have become moribund. As expected, H99-infected mice showed high fungal burdens in the lungs, brains, and kidneys, with the highest fungal burden in the brain (over 10^6^ CFUs) (Figure 5E). The *cbk1*Δ mutant is avirulent in this intravenous infection model as no viable cells were recovered in any organ. Similarly, we could not recover any *sup1* cells from the lungs or the brain, and only detected few fungal cells in the kidney. In contrast, *sup2* suppressor mutants were recovered in all three organs, albeit with reduced fungal burdens (∼10^4^ CFUs in the brain and a few hundred in lungs/kidney) compared to the wildtype H99 control group (Figure 5E). This finding indicates that the *sup2* suppressor, once disseminated into the bloodstream, can invade other organs and replicate. Combined with the earlier observations that 1) both suppressors fully restore growth at host temperature and 2) *sup2* is slightly more CO_2_ tolerant than *sup1*, the observation that only *sup2* can survive, amplify, and persist in animals stresses the importance of CO_2_ tolerance in cryptococcal pathogenesis. Collectively, the results from phagocytosis, the inhalation infection model, and the intravenous infection model, support the hypothesis that failure of the suppressor mutants to disseminate to other organs in the intranasal model is largely due to reduced phagocytosis and inability to escape the lungs. That said, other factors, such as increased systemic clearance by the immune system, could potentially contribute to the containment of the mutant in the lungs. Again, the *cbk1*Δ*ssd1*Δ mutant recapitulated the phenotype of the *sup2* strain in intravenous infection model and other in vitro assays, demonstrating that our observed *sup2* phenotypes are due to disruption of *SSD1*.

## Discussion

Detection of and adaptation to changing CO_2_ levels is an important trait across biological kingdoms and may play a crucial role in the pathogenicity of fungi (Bahn & Mühlschlegel, 2006; Cummins, Selfridge, Sporn, Sznajder, & Taylor, 2014; Hetherington & Raven, 2005). Here we report the identification of genes required for growth at high levels of CO_2_ in the fungal pathogen *C. neoformans*. Multiple pathways important for growth at high temperature, such as the Ras1-Cdc24, CWI, Calcineurin, and RAM pathways, were found to be required for normal growth in high CO_2_, indicating that growth in response to host CO_2_ may be intricately coordinated and co-regulated with response to host temperature. It is therefore likely that both host CO_2_ and host temperature stress hamper related biological functions in cryptococcal cells.

Calcineurin and RAM pathways both appeared as hits in our mutant screen for cryptococcal CO_2_ sensitivity. A previous study found synthetic lethality between the RAM and Calcineurin pathways in *C. neoformans* but not in *S. cerevisiae* (Walton et al., 2006). This corroborates our findings of the key differences between the basidiomycete *C. neoformans* and the ascomycete yeasts. In ascomycete fungal pathogen *C. albicans*, CO_2_ levels are sensed through bicarbonate or cAMP-dependent activation of adenylyl cyclase (Du et al., 2012; Hall et al., 2010). Our results indicate that this pathway does not play a significant role in CO_2_ tolerance in *C. neoformans*. We also found that disruption of the RAM pathway effector kinase Cbk1 caused a severe growth defect at host CO_2_ in *C. neoformans*, but not in *C. albicans*. The vast differences between these organisms in terms of growth response to CO_2_ may reflect the evolutionary distance between these species and/or the distinct niches they normally occupy. Indeed, *C. albicans* is a human commensal and is commonly exposed to host CO_2_ levels. *S. cerevisiae* is a powerful fermenter that thrives in conditions with high levels of CO_2_. For the environmental fungus *C. neoformans*, however, the ability to grow in a CO_2_-enriched condition does not appear to be strongly selected for in the natural environment, and the host level of CO_2_ (∼5% CO_2_) is over 100-fold higher than the ambient air (∼0.04% CO_2_).

The RAM pathway mutants were among the most sensitive mutants to host levels of CO_2_. Remarkably, the growth defects of *cbk1*Δ could be partially restored by single mutations in the genes *PSC1* or *SSD1*. While the PARN ribonuclease-encoding gene *PSC1* represents an uncharacterized protein, *SSD1* is a known suppressor of *cbk1*Δ phenotypes that has been extensively characterized in ascomycete yeasts to regulate the translation of numerous and diverse mRNA transcripts (Hu et al., 2018; Jansen et al., 2009; Lee, Kim, Kang, Yang, & Kim, 2015; L. Li et al., 2009; Wanless et al., 2014). Our genetic interaction analysis indicates that Psc1 likely functions in the same pathway as Ssd1. Interestingly, in *S. cerevisiae*, deletion of *SSD1* can suppress the lethality of *cbk1*Δ but not the cell separation defect, which is regulated by the transcription factor Ace2 (Kurischko, Weiss, Ottey, & Luca, 2005). However, an Ace2 homolog has not been identified in *C. neoformans* or any other basidiomycete (J. Lin et al., 2015). The observation that *cbk1*Δ*psc1*Δ and *cbk1*Δ*ssd1*Δ suppressor mutants partially rescue cell separation defects suggests that *C. neoformans* may primarily utilize Ssd1/Psc1 rather than a potential Ace2 homolog to regulate cell separation. Differential regulation of target mRNA transcripts by Ssd1 and Psc1 may explain the functional divergence of the RAM pathway we observed here between basidiomycete *Cryptococcus* and the ascomycete yeasts. Our observation that the natural suppressors do not restore transcript abundances of CO_2_-associated genes in *cbk1*Δ to WT levels supports a hypothesis that disruption of Ssd1 and Psc1 suppresses *cbk1*Δ mutant’s defects at a post-transcriptional level. *C. neoformans* has been demonstrated to use post-transcriptional regulation to adapt to various host stresses (Bloom et al., 2019; Kalem, Subbiah, Leipheimer, Glazier, & Panepinto, 2021; Stovall, Knowles, Kalem, & Panepinto, 2021). A temperature-sensitive environmental species of *Cryptococcus, C. amylolentus*, fails to initiate host stress-induced translational reprogramming and is non-pathogenic (Bloom et al., 2019). Whether or not translatome reprogramming is initiated in *C. neoformans* in response to host CO_2_, and whether such reprogramming, if occurs, relies on Ssd1 and/or Psc1, has yet to be determined.

## Materials and Methods

### Strains, growth conditions, and microscopy examination

Strains used in this study are listed in Table S2. Unless stated otherwise, all *C. neoformans* cells were maintained at 30°C on yeast peptone dextrose (YPD) media or YPD + CuSO_4_ (25 μM) for strains transformed with P_*CTR4*_-*CBK1*. For morphological examination, all strains were examined under a Zeiss Imager M2 microscope, equipped with an AxioCam MRm camera. For spotting assays, the tested strains were grown overnight in liquid YPD medium at 30°C with shaking at 220 RPM. The cells were then adjusted to the same cell density of OD_600_ = 1 and serially diluted 10-fold. The cell suspensions were then spotted onto YPD agar medium and incubated at the indicated condition for two days. CO_2_ levels were controlled by a VWR CO_2_ incubator or by a Pro-CO_2_ controller (Biospherix, Lacona, NY, USA)

### Genetic manipulation

Gene Deletion Constructs: To delete the gene *SSD1*, a deletion construct with a Nourseothricin (NAT) resistance marker cassette with 5’ and 3’ homology arms to *SSD1* was used. Primers Linlab7974 (gctgcctttgcgtcatctc) and Linlab7976 (ctggccgtcgttttactctcgccttccttctcctta) were used to amplify the 5’ arm from the H99 genome. The 3’ arm was amplified from H99 with primers Linlab7977 (gtcatagctgtttcctgcgattgacattgccgtcttag) and Linlab7979 (cgacctgatcaaactactcgc). The NAT marker was amplified with universal primers M13F and M13R from plasmid pPZP-NATcc. The three pieces were fused together by overlap PCR and amplified with nested primers Linlab7975 (acaatgagccactgccag) and Linlab7977 (tgcgtgttcactactgtagac). To disrupt the gene *PSC1*, a Hygromycin (HYG) marker cassette was used to insert into the PARN domain. To generate the sgRNA for specific targeting to the *SSD1* locus, the *U6* promoter and sgRNA scaffold were amplified from JEC21 genomic DNA and the plasmid pDD162 using primers Linlab7980/Linlab4627 (ttgagtggggtgggtcaattaacagtataccctgccggtg and ggctcaaagagcagatcaatg) and Linlab7981/Linlab4628 (aattgacccaccccactcaagttttagagctagaaatagcaagtt and cctctgacacatgcagctcc). For sgRNA targeted mutation of *PSC1*, the primers Linlab8380/Linlab4627 (tagttgttttcgccgacgccaacagtataccctgccggtg and ggctcaaagagcagatcaatg) were used to amplify the *U6* promoter, and Linlab8381/Linlab4628 (ggcgtcggcgaaaacaactagttttagagctagaaatagcaagtt and cctctgacacatgcagctcc) to amplify the sgRNA scaffold. The *U6* promoter and sgRNA scaffold were fused together by overlap PCR with primers Linlab4594/Linlab4595 (ccatcgatttgcattagaactaaaaacaaagca and ccgctcgagtaaaacaaaaaagcaccgac) to generate the final sgRNA construct as described previously (Fan & Lin, 2018; Jianfeng Lin et al., 2020).

Gene Overexpression Constructs: The *CBK1* overexpression construct was generated by amplifying the *CBK1* open reading frame with primers Linlab7005/BC (ataggccggccatgtcgtatcgcccaatccag) and Linlab7006/BC (cagcatctcgtatcgtcggaag) and cloning the fragment with FseI and PacI into the pXC plasmid backbone (Wang et al., 2012), which contains the promoter of *CTR4* and Neomycin resistance marker. The *CTR4* promoter is highly induced on the copper limiting YPD media. The *MPK1* overexpression construct was generated by amplifying the *MPK1* open reading frame with primers Linlab8326/BC (ataggccggccatggacaatacccctagacac) and Linlab8327/BC (ccttaattaaggctatgataatttctgcctctcc) and cloning the fragment with FseI and AsiSI into a pUC19 plasmid backbone, containing the promoter of *GPD1* and Neomycin resistance marker. The *CDC24* overexpression construct was generated by amplifying the *CDC24* open reading frame with primers Linlab6674/BC (ataggccggccatgtctgtatccggtcccatctc) and Linlab6675/BC (ccttaattaaggataaatctctccttgtggggtacc) and cloning the fragment with FseI and PacI into a pUC19 plasmid backbone, containing the promoter of *CTR4* and Neomycin resistance marker. The overexpression constructs were integrated into the *SH2* locus as described previously (Fan & Lin, 2018; Jianfeng Lin et al., 2020).

Transformation: Constructs for overexpression and deletion were transformed into *Cryptococcus* strains by the TRACE method (Fan & Lin, 2018; Jianfeng Lin et al., 2020), and transformants were selected on YPD medium with 100 μg/mL of nourseothricin (NAT),100 μg/mL of neomycin (NEO), or 200 μg/mL of hygromycin (HYG).

### NanoString RNA profiling

Overnight YPD cultures of H99, *cbk1*Δ, *cbk1*Δ*ssd1*Δ, and *cbk1*Δ*psc1*Δ were washed 2X in PBS and resuspended in RPMI+165mM MOPS, pH 7.4 before quantification on an Invitrogen Countess automated cell counter. Cells were diluted to 7.5×10^5^ cells per mL in 3 mL per well in a 6-well plate. Two wells were used for each biological replicate (n=3) and condition (ambient or 5% CO_2_). Plates were sealed with BreatheEasy sealing membranes (Sigma #Z380059) and incubated in a static incubator at 30°C in ambient air or 5% CO_2_ for 24 hours. Cells were harvested, pelleted at 3,200xg for 5 minutes, and the supernatant was removed. The pellets were then frozen at -80°C and lyophilized overnight. Lyophilized cells were disrupted for 45 seconds with 0.5mm glass beads on an MP Biomedicals FastPrep-24 benchtop homogenizer. RNA was extracted following manufacturer instructions for the Invitrogen PureLink RNA mini-kit with on-column DNAse treatment. Purified RNA was quantified on a NanoDrop OneC spectrophotometer and a total of 100ng per sample was combined with a custom probeset (Dataset S1) from NanoString Technologies according to manufacturer instructions. Probes were hybridized at 65°C for 18 hours, then run on a NanoString nCounter SPRINT profiler according to manufacturer instructions. Data from Reporter Code Count (RCC) files were extracted with nSolver software (version 4.0) and raw counts were exported to Microsoft Excel. Internal negative controls were used to subtract background from raw counts (negative control average + 2 standard deviations). Counts were normalized across samples by total RNA counts. Probes below background were set to a value of 1. Fold change and significance were calculated in Excel after averaging biological triplicates, using a Student t-test (p<0.05). Volcano plot was generated with transformed values (-log[p-value] and log_2_[fold change]) in GraphPad Prism 9. Normalized total counts were used in Morpheus (https://software.broadinstitute.org/morpheus/) to generate a heat map, with hierarchical clustering, one minus Pearson correlation, average linkage method and clustered according to rows and columns.

### Bioinformatics

Whole genome sequencing was performed using the Illumina platform with NovaSeq 6000 at the University of California – Davis Sequencing Center, Novogene USA. A paired-end library with approximately 350 base inserts was constructed for each sample, and all libraries were multiplexed and run in one lane using a read length of 150 bases from either side.

The Illumina reads were first trimmed with Trim Galore v0.6.5 (Krueger, 2021), and then mapped to the *Cryptococcus neoformans* H99 reference genome (FungiDB version 50) using the BWA-MEM algorithm of the BWA aligner v0.7.17(Li, 2013). SAMtools v1.10 (H. Li et al., 2009), Picard Tools v2.16.0 (Broad_Institute), and bcftools v1.13 (Danecek et al., 2021) were used for variant calling from each sample. Variants in the suppressor strains were called with the original *cbk1*Δ mutant as a reference.

The protein diagrams of Psc1 and Ssd1 were made with the illustrator of biological sequences (IBS) software package (Liu et al., 2015).

### Phagocytosis Assays

Mouse macrophage cell line J774A.1 (ATCC^®^ TIB-67™) was acquired from the American Type Culture Collection. Phagocytosis assays were performed using similar procedures as we described previously (J. Lin et al., 2015). Briefly, 1mL of 2×10^5^ J774A.1 macrophages (MΦ) in DMEM media was seeded into a 24 well plate and incubated at 37°C with 5% CO_2_ for 24 hours. *Cryptococcus* strains with a starting OD_600_ of .2 in 3mL of liquid YPD were cultured for 16 hours. Each strain had three technical replicates. The cells were washed three times in sterile H_2_O. 2×10^6^ cryptococcal cells of each strain were opsonized in either 40ul of 100% fetal bovine serum, naïve mouse serum, or mouse serum from LW10 vaccinated A/J mice (Jianfeng Lin et al., 2022; Zhai et al., 2015), for 30 minutes prior to co-incubation with MΦ. Old DMEM media from MΦ was removed and 1mL of fresh DMEM with the opsonized *Cryptococcus* cells were added, followed by a 2-hour incubation at 37°C with 5% CO_2_. The co-culture was then washed six times with warm PBS to remove non-adherent *Cryptococcus* cells. To lyse the macrophages, the cell suspensions were washed with 1mL of cold PBS + 0.01% Triton X. Serial dilutions in PBS of the cell suspensions were then plated onto YNB agar medium and allowed to grow at 30°C for two days to count colony forming units (CFUs).

### Ethical statements

This study was performed according to the guidelines of NIH and the University of Georgia Institutional Animal Care and Use Committee (IACUC). The animal models and procedures used have been approved by the IACUC (AUP protocol numbers: A2017 08-023 and A2020 06-015).

### Murine models of cryptococcosis

Intranasal infection model: Female Balb/C mice of 8-10 weeks old were purchased from the Jackson Labs (Bar Harbor, Maine). Cryptococcal strains were inoculated in 3mL of liquid YPD medium with the initial OD_600_= 0.2 (approximately 10^6^ cell/mL) and incubated for 15 hours at 30°C with shaking. Prior to intranasal infection, cells were washed with sterile saline three times and adjusted to the final concentration of 2×10^5^ cell/mL. Once the mice were sedated with ketamine and xylazine via intraperitoneal injection, 50μL of the cell suspension (1×10^4^ cells per mouse) were inoculated intranasally as previously described (Jianfeng Lin et al., 2022; Zhai, Wu, Wang, Sachs, & Lin, 2012; Zhai et al., 2013; Y. Zhao, Wang, Upadhyay, Xue, & Lin, 2020; Zhu, Zhai, Lin, & Idnurm, 2013). Mice were monitored daily for disease progression. Surviving animals were euthanized at day 35 or 60 post-infection (DPI) and the brain, lungs, and kidneys, were dissected.

Intravenous infection model: Prior to intravenous infections, cryptococcal cells were washed with sterile saline three times and adjusted to the final concentration of 1×10^6^ cell/mL. Mice were sedated with Isoflurane. 100μL of the cell suspension (1×10^5^ cells per mouse) were injected intravenously as previously described (Zhai et al., 2012; Zhai et al., 2013; Y. Zhao et al., 2020; Zhu et al., 2013). After DPI 5, animals were euthanized and the brain, lungs, and kidneys were dissected.

For fungal burden quantifications, dissected organs were homogenized in 2mL of cold sterile PBS using an IKA-T18 homogenizer as we described previously (Zhai et al., 2015; Zhai et al., 2012). Tissue suspensions were serially diluted and plated onto YNB agar medium and incubated at 30°C for 2 days before counting the colony-forming units (CFUs).

## Data Availability

Sequences generated from this research has been deposited to the Sequence Read Archive (SRA) under project accession number: PRJNA791949.

## Acknowledgments

This work was supported by National Institutes of Health (http://www.niaid.nih.gov) (R01AI147541 to D.J.K. and X.L., and R01AI140719 to X.L.). The funder had no role in study design, data collection, and interpretation, or the decision to submit the work for publication. We thank all Lin lab members for their helpful suggestions. We thank Dr. Fanglin Zheng for the plasmid pFZ1, and Dr. Lukasz Kozubowski for the plasmid LKB61.

## Supplementary Information

**Fig. S1.**
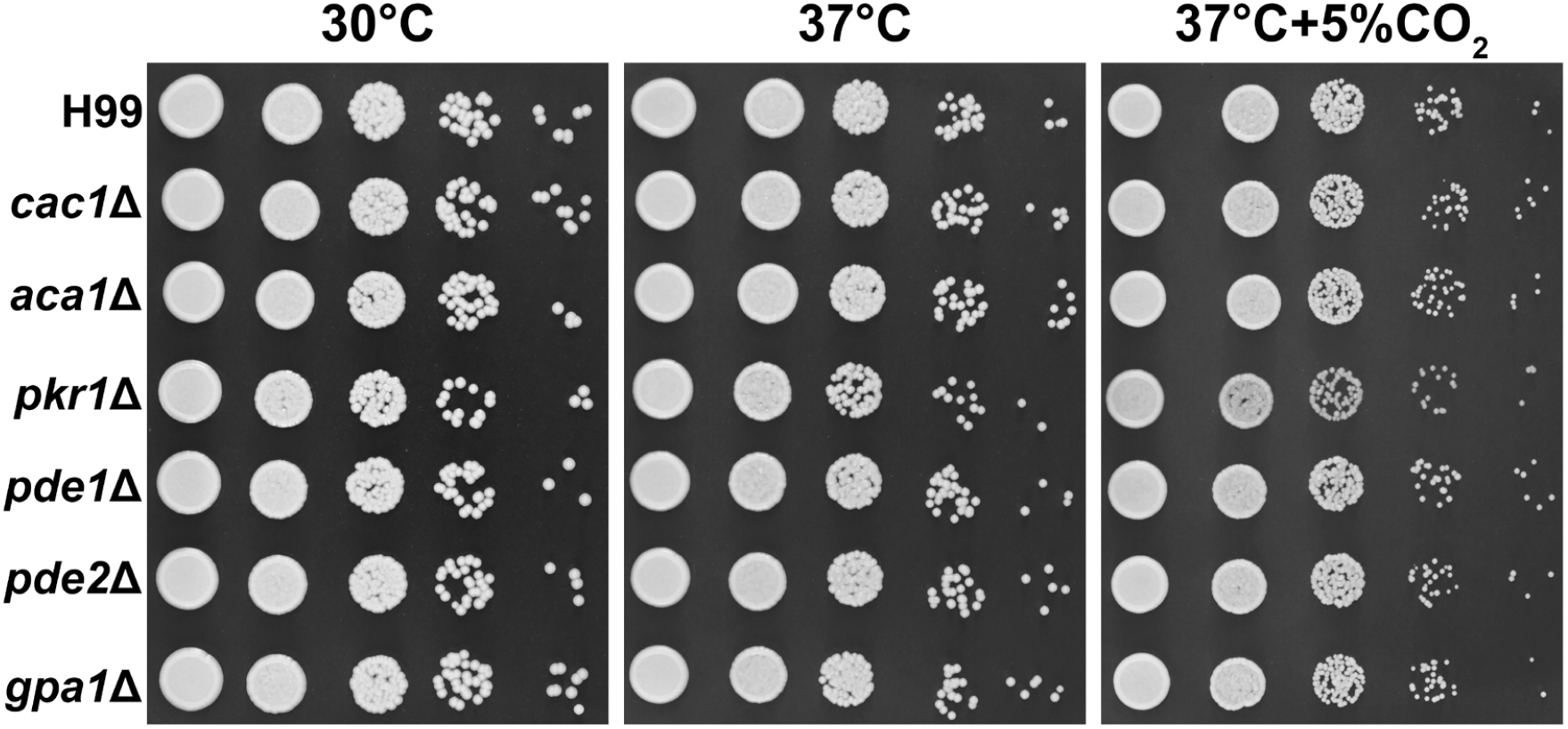
The cAMP pathway is not essential for CO_2_ tolerance. The reference strain H99 and the indicated gene deletion mutants were grown overnight in YPD, serially diluted, and spotted onto solid YPD media plates. Photographs were taken two days after incubation in the indicated condition.

**Fig. S2.**
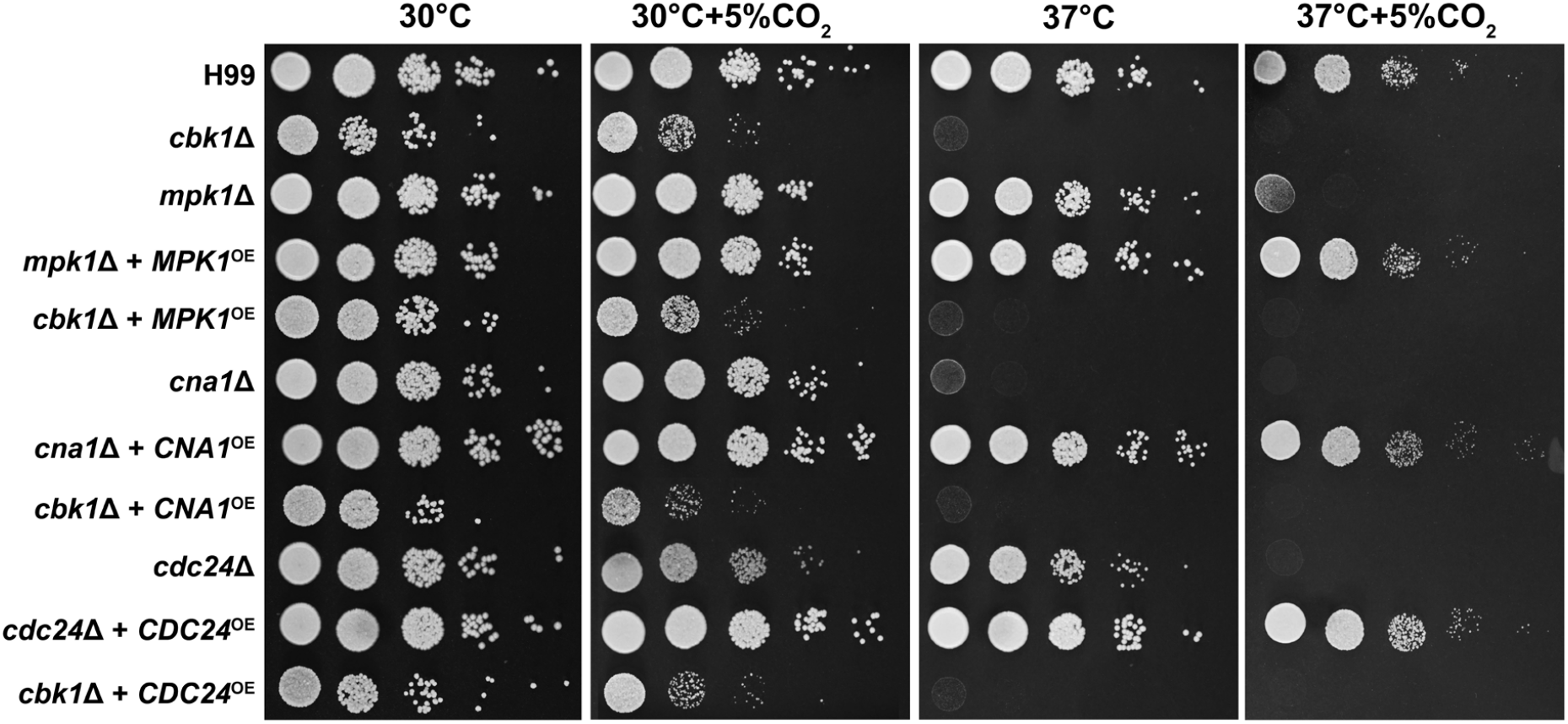
Overexpression of *CDC24, MPK1*, or *CNA1* does not restore growth at host CO_2_ or temperature levels. The strains above were grown overnight in YPD, serially diluted, and spotted onto solid YPD media plates. Photographs were taken two days after incubation in the indicated condition.

**Fig. S3.**
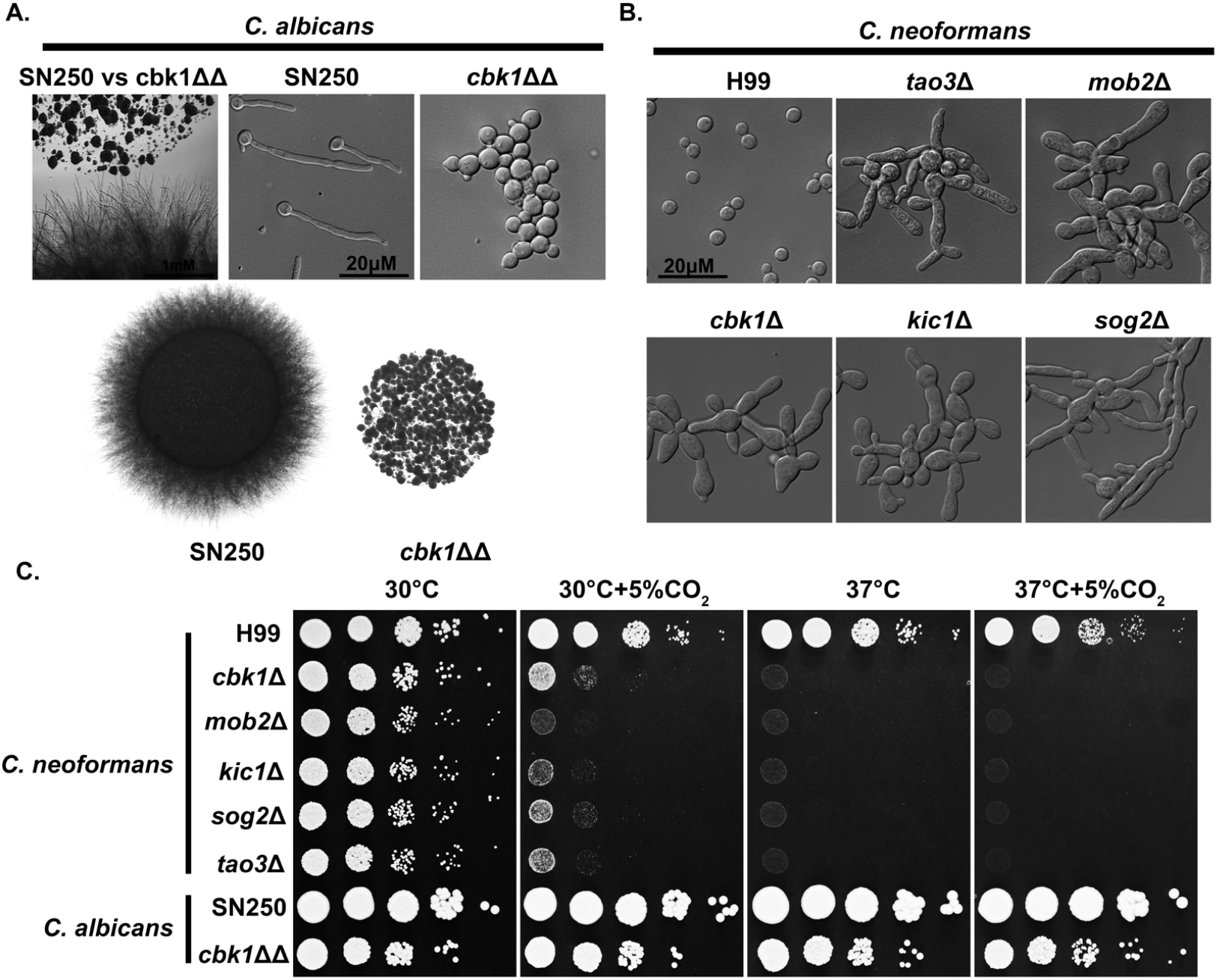
Conserved and divergent roles of the RAM pathway in ascomycete *Candida albicans* and basidiomycete *Cryptococcus neoformans*. (A) The colony and cellular morphology of *Candida albicans* WT strain SN250 and the homozygous cbk1ΔΔ mutant grown on RPMI. (B) The cellular morphology of *Cryptococcus neoformans* WT H99 and RAM pathway mutants cultured in YPD medium. (C) *C. neoformans* and *C. albicans* WT and RAM pathway mutants were serially diluted, spotted onto YPD medium, and incubated for two days at the indicated condition. The image of *C. neoformans* strains is the same as shown in Figure 3A and is included here for direct comparison with *C. albicans* strains.

**Fig. S4.**
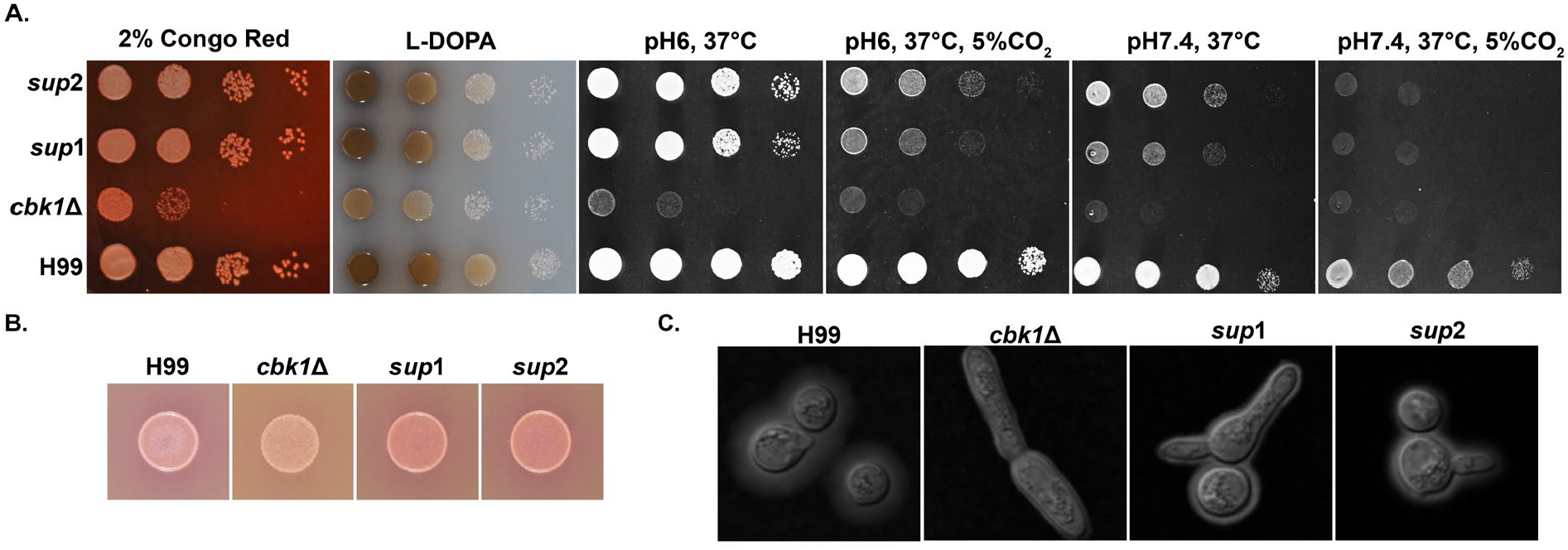
Phenotypic characterization of *cbk1*Δ suppressor mutants. (A, B) The strains above were grown overnight in YPD, serially diluted, and spotted onto the indicated solid media. YPD+2% Congo Red was used to assay cell wall stress tolerance. L-DOPA media was used to assay melanin synthesis ability. RPMI media buffered with MOPS was used to test tolerance to 37°C or 37°C+ 5% CO_2_ at pH6 and pH7.4 to test growth at different pH levels. Photographs were taken two days of incubation. (B) Christensen Urea Agar was used to assay urease activity, indicated by change in media coloration from yellow to pink. (C) The cells incubated for two days on RPMI pH7.4, 37°C+ 5% CO_2_, were stained with India Ink and observed under the microscope to check capsule size.

**Fig. S5.**
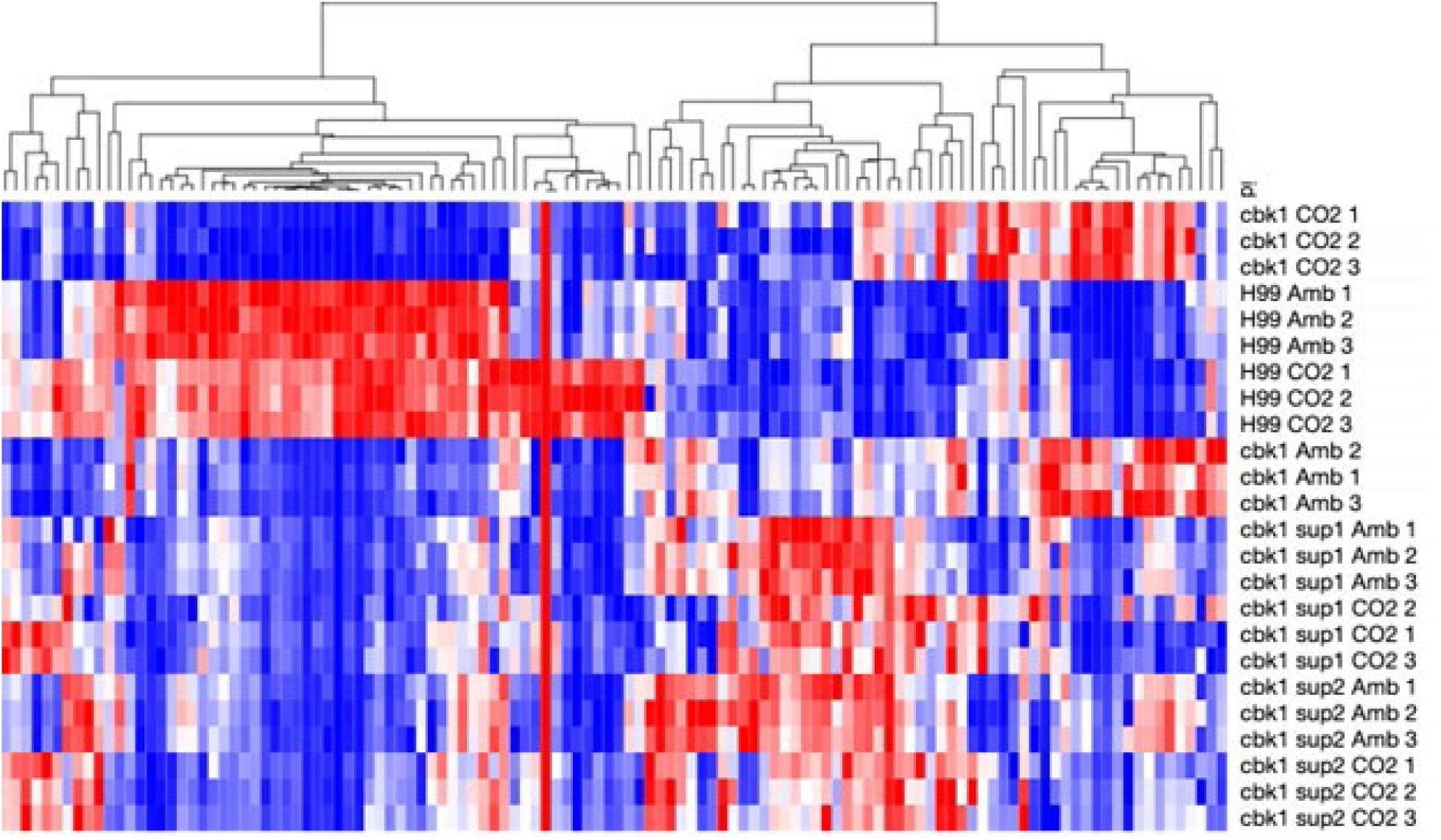
Suppressor mutants do not restore transcript levels of Nanostring targets in *cbk1*Δ. Heatmap showing normalized total RNA counts of NanoString targets in H99, *cbk1*Δ, *sup1*, and *sup2* cultured at either ambient air or 5% CO_2_. Red indicates higher and blue indicates lower transcript abundance.

**Table S1.** Hits from forward genetic screening.

| **Gene ID** | **FungiDB Description** |
| --- | --- |
| CNAG_00097 | myo-inositol transporter, putative |
| CNAG_00293 | Ras-like protein |
| CNAG_00328 | DNA excision repair protein ERCC-5 |
| CNAG_00333 | hypothetical protein |
| CNAG_00405 | ste/ste20/ysk protein kinase |
| CNAG_00414 | Maintenance of killer protein 32, putative |
| CNAG_00443 | hypothetical protein |
| CNAG_00444 | hypothetical protein |
| CNAG_00458 | hypothetical protein |
| CNAG_00503 | hypothetical protein |
| CNAG_00563 | hypothetical protein |
| CNAG_00577 | hypothetical protein |
| CNAG_00699 | transmembrane receptor |
| CNAG_00833 | hypothetical protein |
| CNAG_00888 | calcineurin subunit B |
| CNAG_00968 | hypothetical protein, variant |
| CNAG_00991 | flap endonuclease 1 |
| CNAG_01016 | vacuolar membrane protein |
| CNAG_01038 | hypothetical protein |
| CNAG_01150 | omega-6 fatty acid desaturase (delta-12 desaturase) |
| CNAG_01213 | hypothetical protein |
| CNAG_01255 | hypothetical protein |
| CNAG_01415 | cytoplasmic protein |
| CNAG_01507 | protein CGI121 |
| CNAG_01536 | myosin heavy chain |
| CNAG_01575 | ATP-binding cassette transporter |
| CNAG_01653 | cytokine inducing-glycoprotein |
| CNAG_01845 | AGC/PKC protein kinase |
| CNAG_01864 | hypothetical protein |
| CNAG_01875 | wd-repeat protein |
| CNAG_01918 | cytoskeletal regulatory protein binding protein |
| CNAG_01936 | Sugar transporter |
| CNAG_02232 | RNA polymerase II-associated factor 1 |
| CNAG_02332 | hypothetical protein |
| CNAG_02359 | small subunit ribosomal protein S25e |
| CNAG_02434 | Copper transport protein ATX1 |
| CNAG_02532 | D-amino-acid oxidase |
| CNAG_02586 | Sugar transporter |
| CNAG_02730 | sorting nexin-41 |
| CNAG_03050 | hypothetical protein |
| CNAG_03155 | ENTH domain-containing protein |
| CNAG_03159 | cytoplasmic protein |
| CNAG_03227 | hypothetical protein |
| CNAG_03301 | mitochondrial import inner membrane translocase subunit TIM13 |
| CNAG_03322 | UDP-glucuronate decarboxylase |
| CNAG_03355 | Two-component-like sensor kinase |
| CNAG_03567 | RAM signaling network protein kinase, putative |
| CNAG_03622 | cell polarity protein mor2 |
| CNAG_03634 | DNA-directed RNA polymerase I subunit RPA49 |
| CNAG_03741 | hypothetical protein |
| CNAG_03745 | hypothetical protein |
| CNAG_03824 | solute carrier family 25 (mitochondrial phosphate transporter) |
| CNAG_03918 | ram signaling network protein |
| CNAG_03963 | tyrosine phosphatase |
| CNAG_04048 | 53 kda brg1-associated factor b |
| CNAG_04159 | ariadne-1 |
| CNAG_04243 | cell division control protein 24 |
| CNAG_04351 | methylmalonate-semialdehyde dehydrogenase (acylating) |
| CNAG_04382 | hypothetical protein |
| CNAG_04642 | tetraspanin Tsp2 |
| CNAG_04655 | rab family protein |
| CNAG_04693 | target of rapamycin complex 2 subunit |
| CNAG_04796 | calcineurin a catalytic subunit |
| CNAG_04853 | derlin-2/3 |
| CNAG_04951 | 3-deoxy-7-phosphoheptulonate synthase |
| CNAG_04992 | hypothetical protein |
| CNAG_05021 | hypothetical protein, variant |
| CNAG_05095 | pod-specific dehydrogenase SAC25 |
| CNAG_05114 | peroxisomal copper amine oxidase |
| CNAG_05159 | hypothetical protein |
| CNAG_05299 | oxidoreductase |
| CNAG_05309 | hypothetical protein |
| CNAG_05451 | hypothetical protein |
| CNAG_05604 | hypothetical protein |
| CNAG_05678 | membrane protein |
| CNAG_05789 | hypothetical protein |
| CNAG_05794 | CBK1 kinase activator protein MOB2 |
| CNAG_05882 | class E vacuolar protein-sorting machinery protein HSE1 |
| CNAG_05992 | hypothetical protein |
| CNAG_05998 | rho family protein |
| CNAG_06003 | hypothetical protein |
| CNAG_06218 | Amidase |
| CNAG_06224 | nuclear movement protein nudC |
| CNAG_06373 | mitotic spindle assembly checkpoint protein MAD2B |
| CNAG_06376 | vacuolar membrane protein |
| CNAG_06529 | hypothetical protein |
| CNAG_06583 | hypothetical protein |
| CNAG_06589 | endoribonuclease L-PSP |
| CNAG_06664 | sorting nexin Mvp1 |
| CNAG_06716 | hypothetical protein |
| CNAG_06728 | kinesin |
| CNAG_06796 | serine/arginine repetitive matrix protein 1 |
| CNAG_07358 | hypothetical protein |
| CNAG_07438 | 3-keto sterol reductase |
| CNAG_07448 | urea transporter, putative |
| CNAG_07862 | fumarate reductase |

**Table S2.** Strains and plasmids used.

| **Strain Name** | **Identifier** | **Source** |
| --- | --- | --- |
| FJW9 | H99alpha, *CBK1::NAT<sup>r</sup>* | (1) |
| FJW10 | H99alpha, *MOB2::NAT<sup>r</sup>* | (1) |
| FJW8 | H99alpha, *KIC1::NAT<sup>r</sup>* | (1) |
| AI136 | MATalpha, *TAO3:: NAT<sup>r</sup>* | (1) |
| AI131 | MATalpha, *SOG2:: NAT<sup>r</sup>* | (1) |
| SN250 | HIS-, LEU-, ARG- | (2) |
| $\Delta\Delta$cbk1 | SN250, $\Delta\Delta$cbk1: *HIS-, LEU-, ARG-* | (2) |
| *cna1*Δ | H99alpha *CNA1::NAT<sup>r</sup>* | FGSC deletion set<br>Plate 46 Well E12 |
| *cdc24*Δ | H99alpha, *CDC24::NAT<sup>r</sup>* | FGSC deletion set<br>Plate 33 Well C4 |
| *mpk1*Δ | H99alpha, *MPK1::NAT<sup>r</sup>* | FGSC deletion set<br>Plate 11 Well A5 |
| BC1449 | H99alpha, *P<sub>CTR4</sub>-CBK1-mCherry-NEO<sup>r</sup>* | This study. |
| BC1281 | H99alpha, *CDC24::NAT<sup>r</sup>*, *P<sub>CTR4</sub>-CBK1-mCherry- NEO<sup>r</sup>* | This study. |
| BC1282 | H99alpha, *CDC24::NAT<sup>r</sup>*, *P<sub>CTR4</sub>-CBK1-mCherry- NEO<sup>r</sup>* | This study. |
| BC1283 | H99alpha, *CNA1::NAT<sup>r</sup>*, *P<sub>CTR4</sub>-CBK1-mCherry- NEO<sup>r</sup>* | This study. |
| BC1284 | H99alpha, *CNA1::NAT<sup>r</sup>*, *P<sub>CTR4</sub>-CBK1-mCherry- NEO<sup>r</sup>* | This study. |
| BC1285 | H99alpha, *MPK1::NAT<sup>r</sup>*, *P<sub>CTR4</sub>-CBK1-mCherry- NEO<sup>r</sup>* | This study. |
| BC1286 | H99alpha, *MPK1::NAT<sup>r</sup>*, *P<sub>CTR4</sub>-CBK1-mCherry- NEO<sup>r</sup>* | This study. |
| BC650 | H99alpha, *CDC24::NAT<sup>r</sup>*, *P<sub>CTR4</sub>-mNeonGreen-CDC24-NEO<sup>r</sup>* | This study. |
| BC669 | H99alpha, *CBK1::NAT<sup>r</sup>*, *P<sub>CTR4</sub>-CBK1-mCherry- NEO<sup>r</sup>* | This study. |
| BC1356 | H99alpha, *MPK1::NAT<sup>r</sup>*, *P<sub>GPD1</sub>-mNeonGreen-MPK1-NEO<sup>r</sup>* | This study. |
| LK214 | H99a, *CNA1::NEO<sup>r</sup>*, *P<sub>H3</sub>-GFP-CNA1-NAT<sup>r</sup>* | (3) |
| BC1357 | H99alpha, *CBK1::NAT<sup>r</sup>*, *P<sub>CTR4</sub>-mNeonGreen-CDC24-NEO<sup>r</sup>* | This study. |
| BC1358 | H99alpha, *CBK1::NAT<sup>r</sup>*, P<sub>GPD1</sub>-mNeonGreen-*MPK1-NEO<sup>r</sup>* | This study. |
| BC1359 | H99alpha, *CNA1::NAT<sup>r</sup>*, P<sub>GPD1</sub>-mCherry-*CNA1-HYG<sup>r</sup>* | This study. |
| BC1068 | H99alpha, *CBK1::NAT<sup>r</sup>*, SUP1 | This study. |
| BC1076 | H99alpha, *CBK1::NAT<sup>r</sup>*, SUP2 | This study. |
| BC1239 | H99alpha, *CBK1::NAT<sup>r</sup>*, *SSD1::NEO<sup>r</sup>* | This study. |
| BC1241 | H99alpha, *SSD1::NEO<sup>r</sup>* | This study. |
| YSB42 | H99alpha, *CAC1::NAT<sup>r</sup>* STM#159 | (4) |
| *aca1Δ* | H99alpha, *ACA1::NAT<sup>r</sup>* | FGSC deletion set<br>Plate 32 Well H6 |
| *pkr1Δ* | H99alpha, *PKR1::NAT<sup>r</sup>* | FGSC deletion set<br>Plate 33 Well H7 |
| *pde1Δ* | H99alpha, *PDR1::NAT<sup>r</sup>* | FGSC deletion set<br>Plate 10 Well A12 |
| *pde2Δ* | H99alpha, *PDR2::NAT<sup>r</sup>* | FGSC deletion set<br>Plate 11 Well H7 |
| YSB83 | H99alpha, *GPA1::NAT<sup>r</sup>* STM#5 | (5) |
| BZ36 | H99alpha, *CLN1::NAT<sup>r</sup>* | This study. |
| *ecm2201Δ* | H99alpha, *ECM2201::NAT<sup>r</sup>* | FGSC deletion set<br>Plate 2 Well A10 |
| *kin4Δ* | H99alpha, *KIN4::NAT<sup>r</sup>* | Bahn Kinase Deletion<br>Set Plate 4 Well G8<br>(6) |
| **Plasmid Name** | **Identifier** | **Source** |
| pPZP-NATcc | pPZP-NATcc | (7) |
| pDD162 | pDD162 | (8) |
| pFZ1-*CDC24* | P<sub>CTR4-2</sub>-mNeonGreen-*CDC24*(H99)-*NEO<sup>r</sup>* | This study. |
| pXC-*CBK1*-mCh | P<sub>CTR4-2</sub>-*CDC24*(H99)-mCherry-*NEO<sup>r</sup>* | This study. |
| LKB61 | P<sub>GPD1</sub>-mCherry-*CNA1-HYG<sup>r</sup>* | (3) |
| pUC19-*MPK1*-mNG | P<sub>GPD1</sub>-*MPK1*-mNeonGreen-*NEO<sup>r</sup>* | This study. |

